# hcga: Highly Comparative Graph Analysis for network phenotyping

**DOI:** 10.1101/2020.09.25.312926

**Authors:** Robert L. Peach, Alexis Arnaudon, Julia A. Schmidt, Henry A. Palasciano, Nathan R. Bernier, Kim Jelfs, Sophia Yaliraki, Mauricio Barahona

**Author notes:** Blue Brain Project, École polytechnique fédérale de Lausanne (EPFL), Campus Biotech, 1202 Geneva, Switzerland.

## Abstract

Networks are widely used as mathematical models of complex systems across many scientific disciplines, not only in biology and medicine but also in the social sciences, physics, computing and engineering. Decades of work have produced a vast corpus of research characterising the topological, combinatorial, statistical and spectral properties of graphs. Each graph property can be thought of as a feature that captures important (and some times overlapping) characteristics of a network. In the analysis of real-world graphs, it is crucial to integrate systematically a large number of diverse graph features in order to characterise and classify networks, as well as to aid network-based scientific discovery. In this paper, we introduce HCGA, a framework for highly comparative analysis of graph data sets that computes several thousands of graph features from any given network. HCGA also offers a suite of statistical learning and data analysis tools for automated identification and selection of important and interpretable features underpinning the characterisation of graph data sets. We show that HCGA outperforms other methodologies on supervised classification tasks on benchmark data sets whilst retaining the interpretability of network features. We also illustrate how HCGA can be used for network-based discovery through two examples where data is naturally represented as graphs: the clustering of a data set of images of neuronal morphologies, and a regression problem to predict charge transfer in organic semiconductors based on their structure. HCGA is an open platform that can be expanded to include further graph properties and statistical learning tools to allow researchers to leverage the wide breadth of graph-theoretical research to quantitatively analyse and draw insights from network data.

## 1 Introduction

Graphs provide an elegant and powerful formalism to represent complex systems [1]. Across many scientific disciplines there are increasing volumes of data that are naturally described as graphs (or networks) and can leverage the extensive results in graph theory to solve research problems. Among many others, examples include linking the structural motifs of proteins and their function [2, 3, 4], aiding the diagnosis of diseases using fMRI data [5], understanding structural properties of organic crystal structures for electron transport [6], or modelling network flows, e.g., city traffic [7], information (or misinformation) spread in a social network [8, 9], or topic affinity in a citation network [10]. The growing importance of such network data has driven the development of a multitude of methods for investigating and revealing relevant topological, combinatorial, statistical and spectral properties of graphs, e.g., node centralities [11, 12], assortativity [13, 14], path-based properties [15], graph distance measures [16, 17], connectivity [18], or community detection [19, 20], to name but a few in the highly interdisciplinary area of network science.

When analysing network data sets, or when posing a research question in terms of network properties, it is usually not immediately clear which graph-theoretical methodologies should be used; particularly in real-world applications, which do not conform to clear-cut constructive assumptions and, hence, where differences between complex systems may be subtle. For example, we may want to identify characteristics of a social network that aid the spread of fake news [21], or identify structural properties that would help us predict the toxicity of a given molecule [22] without restricting *a priori* the types of graph properties to be considered. The viability and enormous potential of highly-comparative data-driven analyses of graphs was previously shown [23], however, existing software packages that derive summary statistics of graphs often only compute a small set of features that are usually chosen to target a subject area (e.g., neuroscience [24] or biology [25]). Despite the myriad of graph properties, there currently exists no systematic way to leverage the wide spectrum of available measures to identify graph features that best charaterise a given problem.

Here, we introduce Highly Comparative Graph Analysis (HCGA), a modular, expandable Python software package ^2^ that allows researchers to perform massive graph feature extraction together with statistical learning and analysis of feature importance. Our computational framework is illustrated in Figure 1 and takes both inspiration and ideas from HCTSA and CATCH22, powerful frameworks for time-series feature extraction [26, 27, 28, 29]. Given a set of complex systems modelled as networks representing, e.g., molecules, proteins, neuronal morphologies, transportation routes, ecological or social networks, HCGA first extracts from each network in the set a few thousand graph features, each encoding a different interpretable network property. The selection of features to be extracted is flexible and can be adapted by the researcher to the particular problem at hand. Following the feature extraction step, each graph in the data set is described as a high-dimensional feature vector, and the whole data set is encoded as a feature matrix. To facilitate data-driven analyses of the data set, HCGA includes a suite of tools for statistical learning aimed at classification, regression, and unsupervised learning. Since HCGA preserves the interpretability of the features, our framework also includes in-depth feature importance analysis using Shapley Additive Values (SHAP) to aid feature selection aimed at deriving scientific insights [30]. HCGA thus removes the time-consuming and subjective task of implementing individual graph-theoretical methods for the analysis of network data sets. The structure of HCGA is modular and open-source, allowing researchers from any research area to contribute further graph-theoretical features; to extend the statistical learning tools; or to improve visualisation modules for the use of the research community.

**Figure 1:**
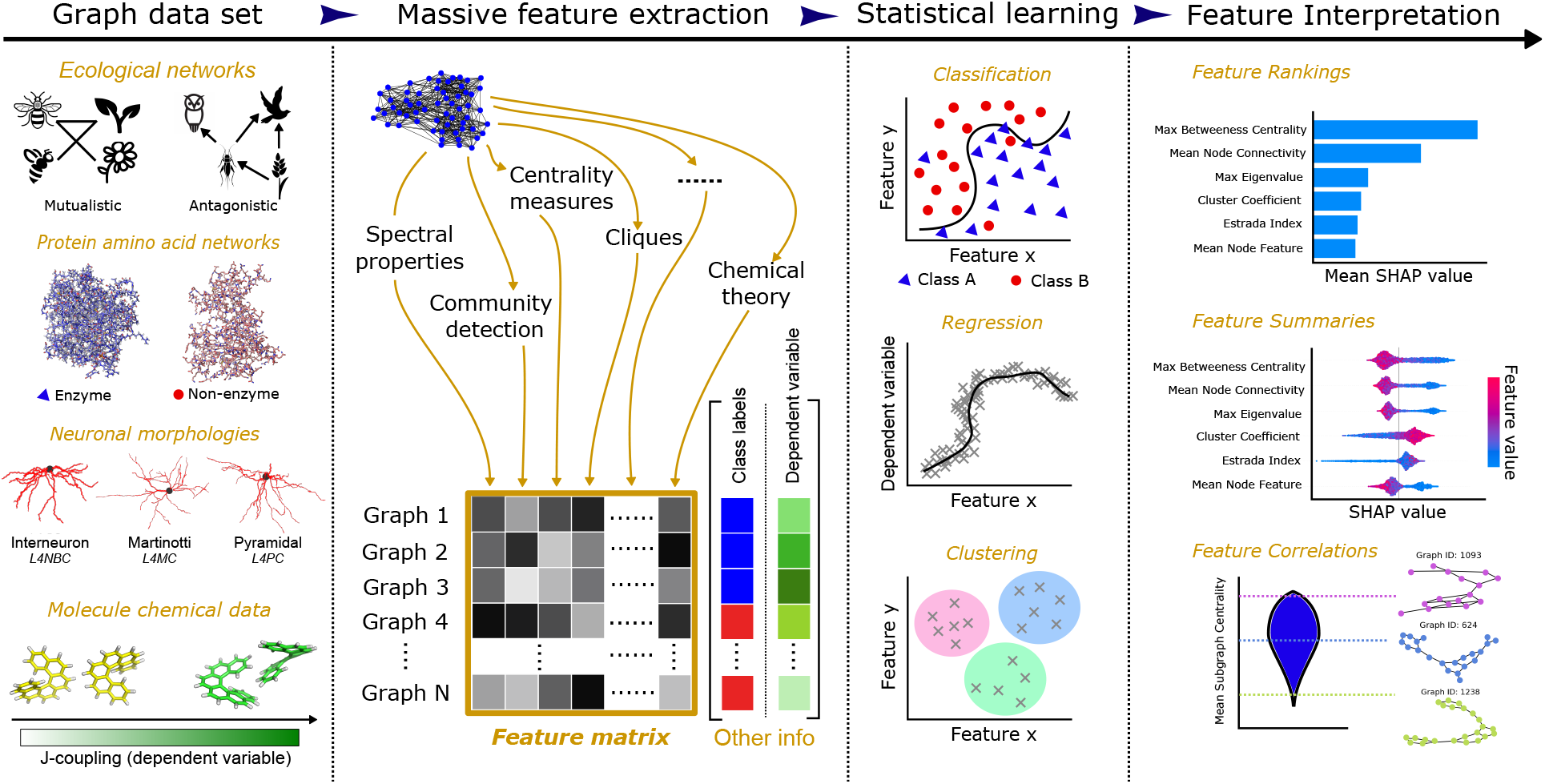
Overview of the HCGA pipeline. HCGA can be applied to a wide range of graph data sets, such as ensembles of synthetic graphs, molecular structures, cell morphologies, social and ecological networks, among many others. HCGA first constructs a large feature matrix for the data set by computing for each network a wide range of graph-theoretical properties (several thousands) that have been compiled from classic and more recent literature. This feature matrix is then used to perform statistical analyses on the graph data set, including supervised and unsupervised learning tasks (e.g., classification, clustering, regression) in conjunction with any additional information known, such as class labels, a continuous dependent variable, or node features. HCGA preserves the interpretability of the features and provides tools (e.g., Shapley values) to characterise the importance of features for the prediction task as an aid to get insights into network-based discovery for the data set at hand. HCGA is an open platform which allows for the expansion of the set of graph-theoretical features, as well as allowing for the inclusion of additional statistical analyses.

**Figure 2:**
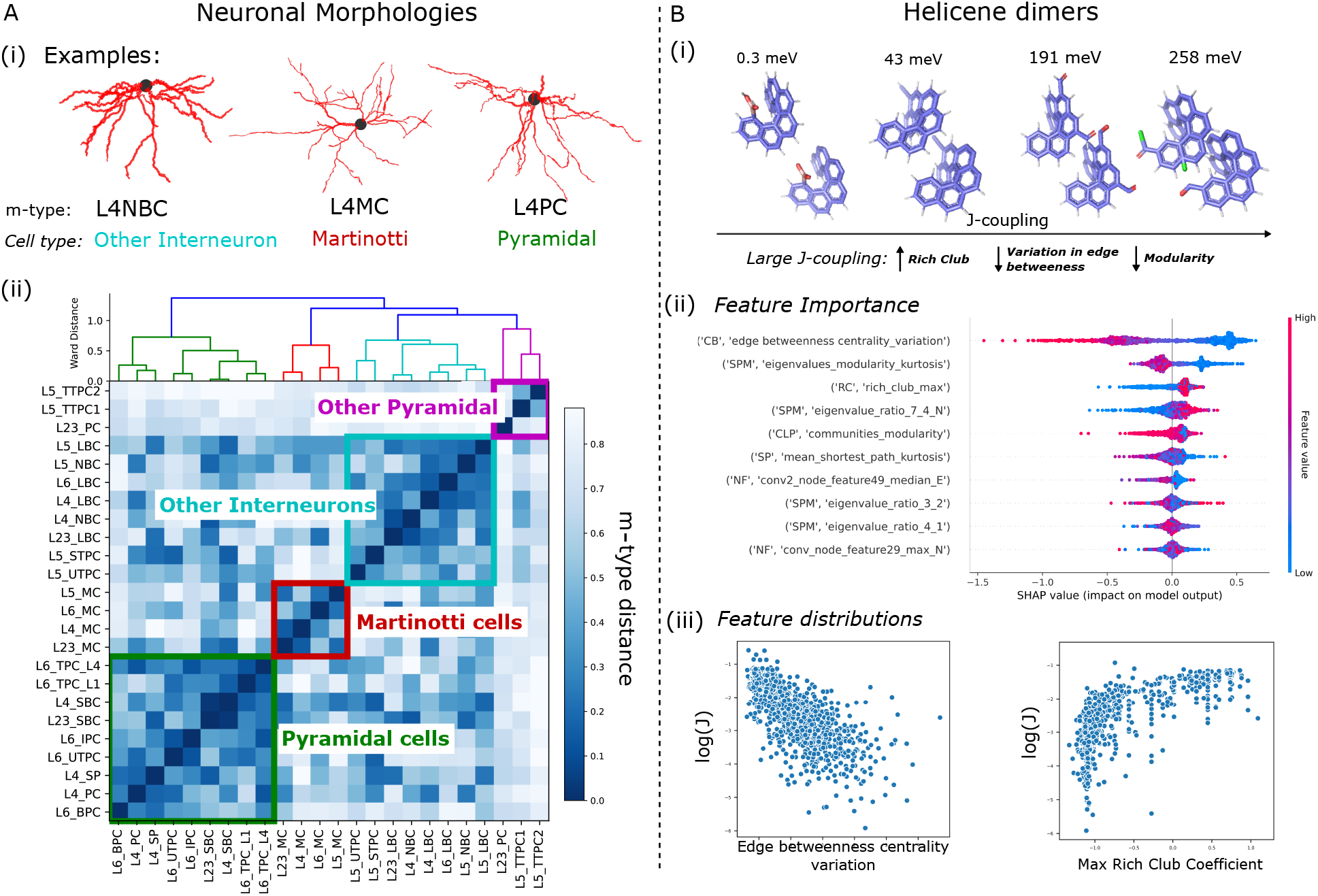
HCGA recovers biological clusters of neuronal morphologies and uncovers interpretable structural properties for effective electronic coupling in semiconductor crystals. (A) *Neuronal morphologies*. (i) Three examples of neuron morphologies for three different cell types which can be further distinguished by the cortical layer from which they were extracted (in this case from L4). (ii) The inverse classification accuracies between m-type pairs using the HCGA features was used to produce a similarity matrix. Clustering of this similarity matrix recovers the three main morphological cell types (Pyramidal, Martinotti and interneurons) and reveals a new cluster consisting of the most common pyramidal cells. Finer clusters group pairs or triplets of the same m-types from different layers. (B) *Helicene dimers*. Four examples of helicene dimers at different magnitudes of J-coupling, coloured by atom-type. (ii) The SHAP value for each individual sample for the top 10 features listed in descending order. Each sample is coloured by their relative feature value normalised between 0 and 1. The sum of absolute SHAP values for individual samples defines the total feature importance SHAP value. (iii) The relationship between log(*J*) and the top two features are illustrated in a scatter diagram; we notice a strong negative trend with the variation of edge betweenness centrality and a positive non-linear trend with the maximum rich club coefficient.

## 2 Results

We illustrate how HCGA can be employed in three tasks aimed at network-based scientific discovery: supervised classification, regression, and unsupervised clustering.

### 2.1 Supervised classification of benchmark graph data sets from biochemistry and social science

We evaluated the performance of HCGA for supervised classification using known collections of labelled graphs that have been used as benchmarks. The sets comprise five biochemical data sets (proteins, enzymes, d&d, NCI1, mutag) and six standard social media data sets (collab, reddit-binary, reddit-multi-5k, reddit-multi-12k, IMDB-binary, IMDB-multi); see Table 2 for details. In Tables 3 and 4, we show the classification accuracy achieved on the biochemical and social data sets, respectively, using an XGBoost classifier (with default settings) applied to the top 100 features extracted with HCGA. Our results show that HCGA achieves top accuracy without optimising the hyperparameters of the XGBoost classifier (as would be the case in a realistic user case scenario) when compared to popular deep-learning methodologies and Kernel algorithms under the Fair Comparison protocol [31]. If we choose the best subset of features through cross-validation, the results are improved further (still without optimising the classifier).

**Table 1:**
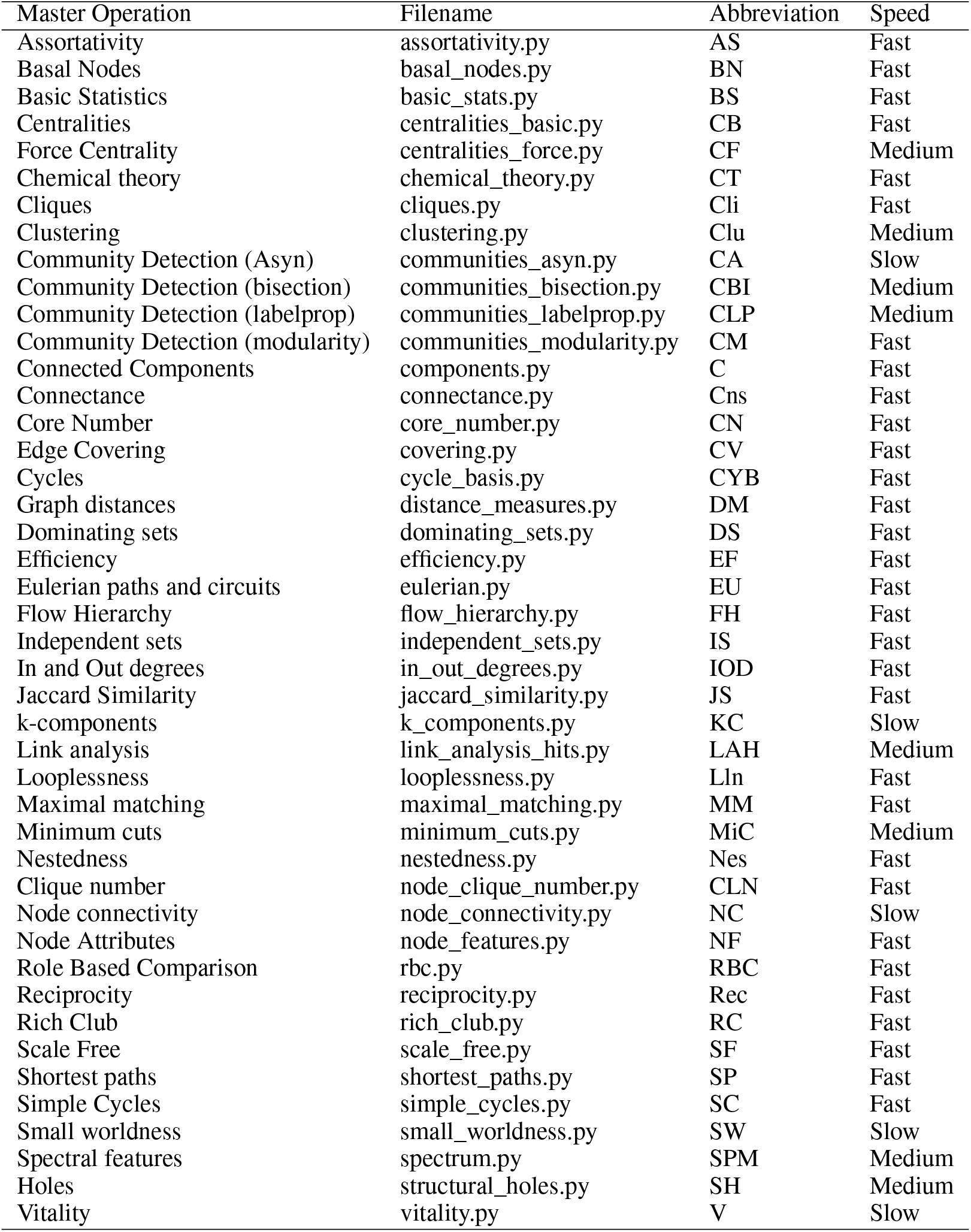
List of master operations and the associated filename and shortname in the HCGA package at time of publication. The speed indicates the assigned computation speed for a given master operation. The speed labels are assigned manually after thorough testing on various graph types. Users can choose to ignore features that are slow to compute, providing flexibility to the user (see A.3.2).

**Table 2:**
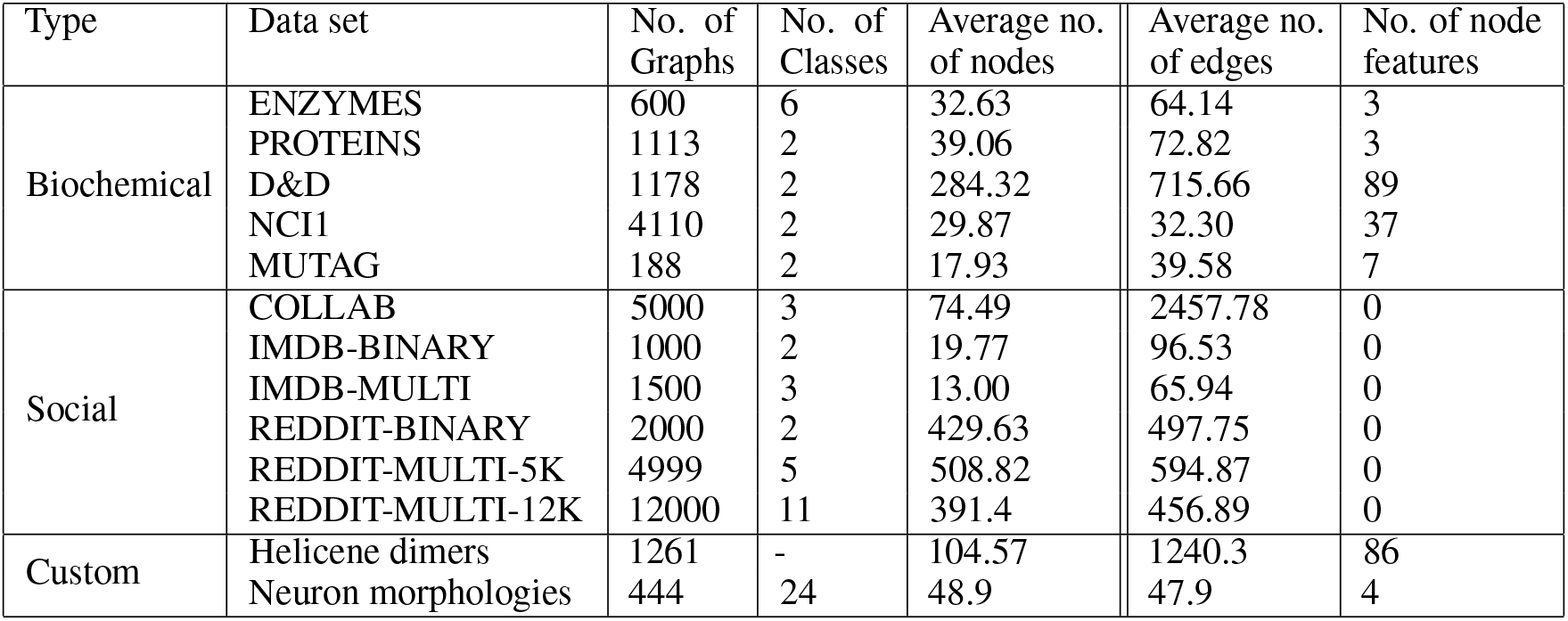
Statistics of the data sets. When node features are present we include these in the classification for comparability with existing methodologies.

**Table 3:**
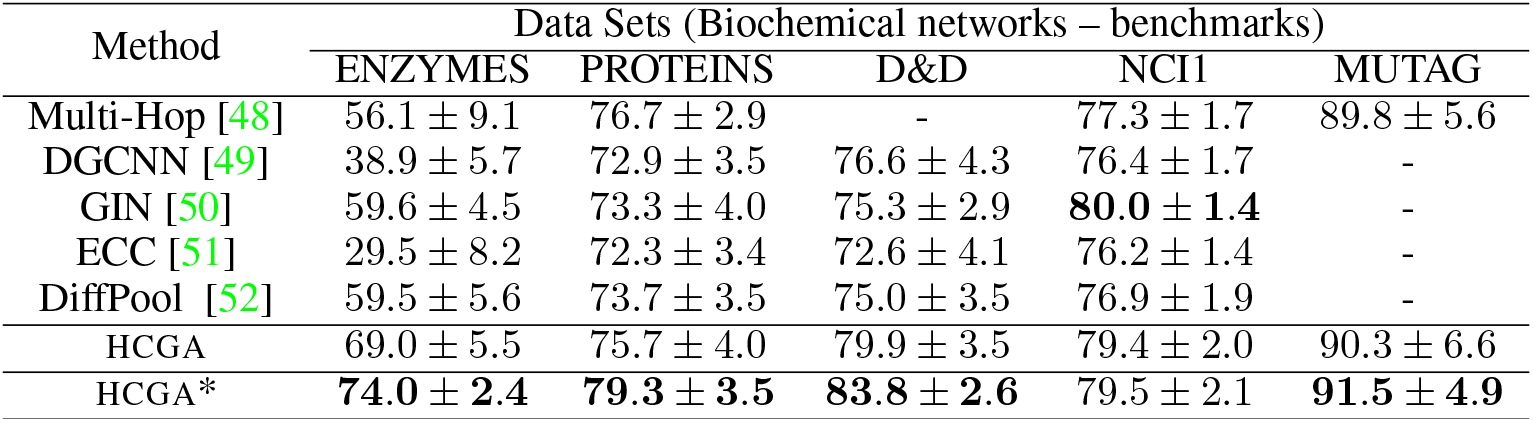
Classification results on biochemical benchmark data sets obtained by HCGA using the XGBoost classifier and compared against other methodologies. The HCGA accuracies are obtained from the top 100 features, whereas HCGA∗ accuracies are obtained from features selected by optimising against a validation set. The hyperparameters of the XGBoost classifier were not optimised for any of the HCGA runs. The classification accuracies for MUTAG are not available for most methods due to long computation times; however we retain this example to show that HCGA is also capable of analysing large data sets.

**Table 4:**
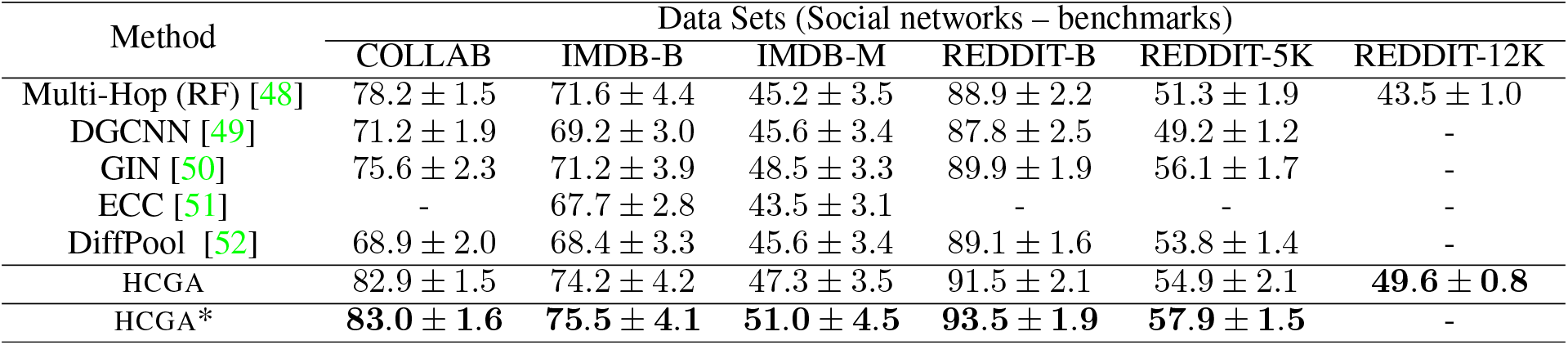
Classification results on social benchmark data sets obtained by HCGA using the XGBoost classifier and compared against other methodologies as described in Table 3. The classification accuracies for REDDIT-12K are not available for most methods due to long computation times; however we retain this example to show that HCGA is also capable of analysing large data sets.

| Method | Data Sets (Social networks – benchmarks) |  |  |  |  |  |
| --- | --- | --- | --- | --- | --- | --- |
|  | COLLAB | IMDB-B | IMDB-M | REDDIT-B | REDDIT-5K | REDDIT-12K |
| Multi-Hop (RF) [48] | $78.2 \pm 1.5$ | $71.6 \pm 4.4$ | $45.2 \pm 3.5$ | $88.9 \pm 2.2$ | $51.3 \pm 1.9$ | $43.5 \pm 1.0$ |
| DGCNN [49] | $71.2 \pm 1.9$ | $69.2 \pm 3.0$ | $45.6 \pm 3.4$ | $87.8 \pm 2.5$ | $49.2 \pm 1.2$ | - |
| GIN [50] | $75.6 \pm 2.3$ | $71.2 \pm 3.9$ | $48.5 \pm 3.3$ | $89.9 \pm 1.9$ | $56.1 \pm 1.7$ | - |
| ECC [51] | - | $67.7 \pm 2.8$ | $43.5 \pm 3.1$ | - | - | - |
| DiffPool [52] | $68.9 \pm 2.0$ | $68.4 \pm 3.3$ | $45.6 \pm 3.4$ | $89.1 \pm 1.6$ | $53.8 \pm 1.4$ | - |
| HCGA | $82.9 \pm 1.5$ | $74.2 \pm 4.2$ | $47.3 \pm 3.5$ | $91.5 \pm 2.1$ | $54.9 \pm 2.1$ | **$49.6 \pm 0.8$** |
| HCGA* | **$83.0 \pm 1.6$** | **$75.5 \pm 4.1$** | **$51.0 \pm 4.5$** | **$93.5 \pm 1.9$** | **$57.9 \pm 1.5$** | - |

In addition to its high classification performance, HCGA preserves the interpretability of features, which can then be ordered according to their importance for each modelling problem. In STAR Methods C.2 we exemplify a more detailed analysis of the features underpinning the classification of the benchmark proteins data set (classifying proteins as enzymatic or non-enzymatic). Among the features that display the largest impact on the prediction task we find an increased number of cliques in enzymatic proteins, reflecting the structurally stable modules necessary for catalysis (Figure 3). Our deep-dive analysis also reveals that, despite being derived from seemingly different areas of graph theory, many top features are highly correlated (Figure 4) and provide similar predictive power. This observation reflects the mathematical relationships, sometimes not explicitly recognised, between many graph features, which can still afford complementary descriptions of the data. These results highlight the need to consider a broad range of features in the analysis of networks, rather than relying on a narrow set of properties.

**Figure 3:**
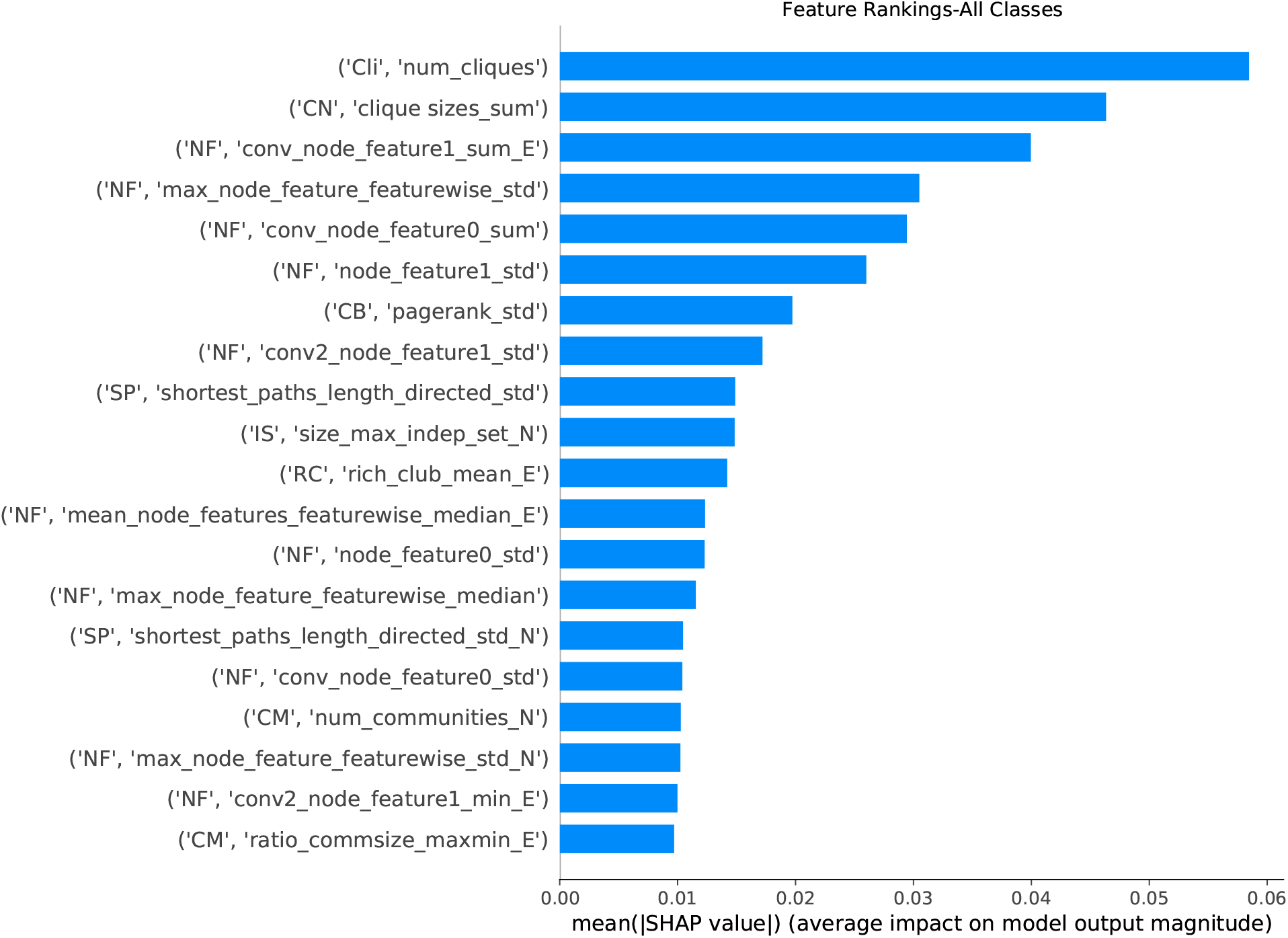
Top 20 features by SHAP score for PROTEINS data set. The features are sorted in descending order by importance on model prediction. The features are indicated by their class ID and their feature name.

**Figure 4:**
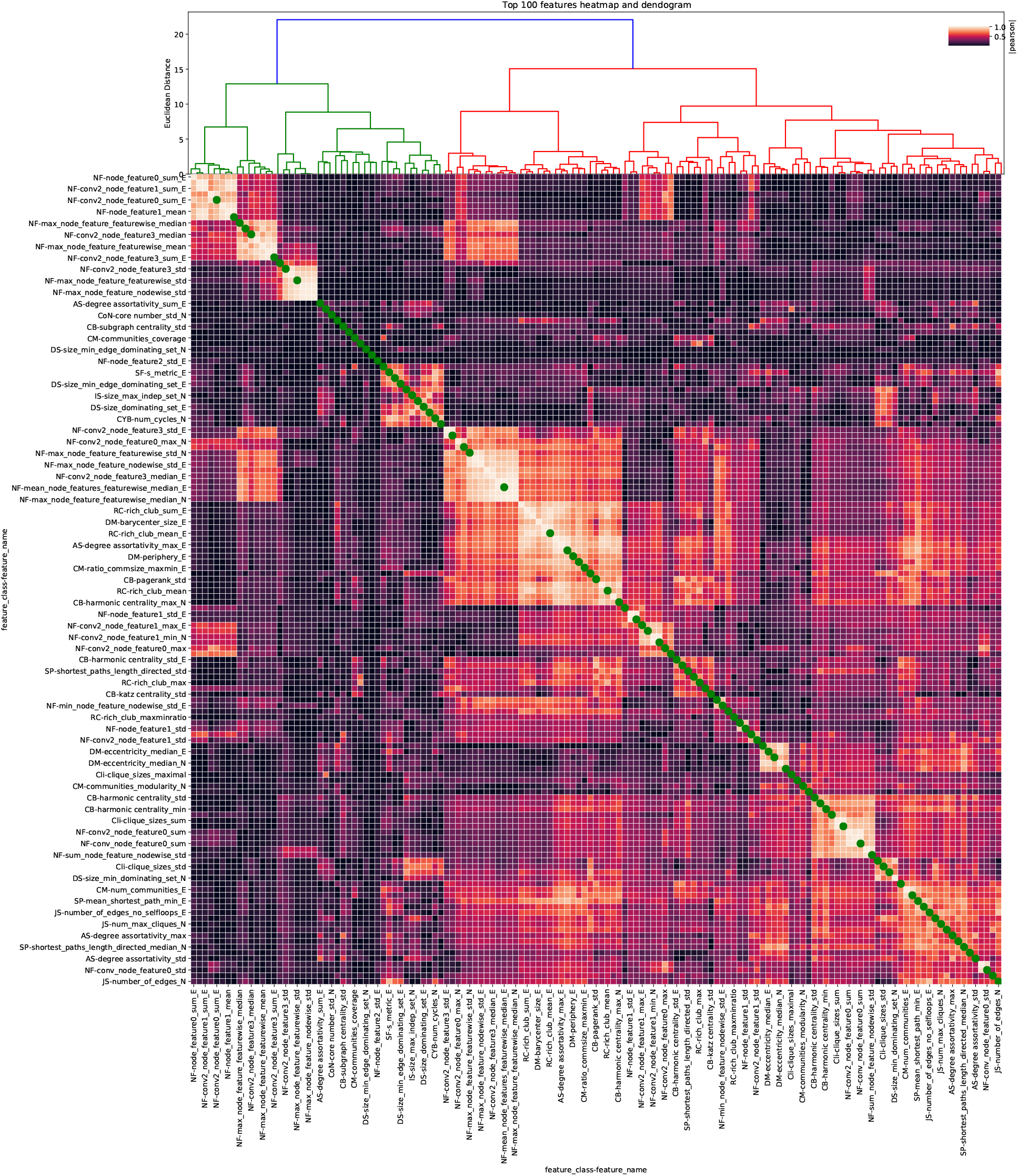
Clustered correlation matrix of features for proteins data set. The absolute correlation coefficient between the top 70 features is computed and collected in a correlation matrix. The dendogram (top) corresponds to a clustering of the correlation matrix using the Ward hierarchical linkage algorithm. Green dots indicate features that were selected after reducing the feature set to 100 minimally correlated features (not all selected features are shown here).

### 2.2 Unsupervised clustering: mapping morphological neuron types onto cell types based on network features

To go beyond supervised classification of benchmark data sets, we used HCGA to analyse a data set of 444 neuronal cells from 6 layers (L1 to L6) in the rat somatosensory cortex belonging to 24 different morphological types (m-types) [32, 33]. The neurons are also labelled independently as belonging to three different cell types: pyramidal neurons, interneurons, and Martinotti cells.

We represent the morphology of each neuron as a rooted tree, with the soma as the root and basal dendrites and axons as branches extending from the soma (Figure 2A(i)). A graph is then constructed for each neuron by assigning a node to each section between two branching points of the morphology. Each node is also assigned features: path length, mean diameter of each branch, and an annotation to differentiate the soma from other nodes. See STAR Methods A.2.3 for more details.

Using HCGA, we extracted 2112 features from the graph of each neuron. Our aim is to compare the similarities between neuron morphological types and their correspondence with cell types. To do so, we construct a similarity matrix between m-types by computing the mean 10-fold classification accuracy between each pair of m-types: low classification accuracy between m-types indicates high pairwise similarity (Figure 2A(ii) and STAR Methods A.2.3). Applying a simple hierarchical clustering with Ward’s linkage to this similarity matrix, we find clusters of m-types that map well to cell types. In particular, a robust clustering into four clusters naturally recovers biologically meaningful groupings of m-types into cell types: pyramidal, Martinotti, other interneurons, as well as a group of the three most common pyramidal m-types (L5_TTPC1, L5_TTPC2 and L23_PC). At a finer resolution, we note that clusters consist of pairs or triplets of the same m-types belonging to different cortical layers. A deeper analysis of feature importance in this data set may reveal further morphological characteristics that define cell types across layers, but such detailed study is beyond the scope of this initial illustrative analysis.

### 2.3 Regressing graph features of helicene structures against their electronic transport

As a second example of a task in scientific discovery, we apply HCGA to predict the electronic properties of helicenes from their structure. Helicenes are graphene-type spiral molecules with promising optical and electronic applications due to their axial chirality [34]. We use a recently curated data set composed of 1344 helicene dimers (i.e., pairs of helicenes in the same translational-motif orientation) [6], see Figure 2B(i). For each helicene dimer, we create a simple molecular graph representing atoms and their van der Waals interactions: an edge is present if the separation between two atoms is lower than the sum of their Van der Waals radii (plus a buffer of 2Å), and the edge weight is given by the inverse distance (more details are given in STAR Methods A.2.4). The atom types are included as one-hot encoded node features. For each dimer, we aimed to predict the electronic transfer integral *J*, or *J*-coupling, which describes the ease with which a charge carrier (electron or hole) can hop from one molecule to another. The *J*-couplings for each helicene dimer were previously calculated using a hybrid DFT molecular pair calculation and the projective method [35] (see STAR Methods A.2.4). The *J*-coupling depends on various factors such as the frontier orbitals of the molecule, the molecular packing (i.e., the relative orientations and distances between molecules [36]), as well as molecular vibrations.

Since the *J*-coupling is a continuous variable, we treat this problem as a regression task. We used HCGA to extract 2531 features from each molecular graph and regressed them against the logarithm of the transfer integral *log*(*J*). Using the full set of features, we achieve a mean absolute error (MAE) of 0.355 ± 0.022 on our evaluation set. To put this into perspective, it is generally expected that the shorter the distance between molecules the higher the charge transfer between them. However, if we use the distance between molecules in the dimer as the only feature, we obtain an MAE of 1.58±, an almost five-fold decrease in accuracy with respect to the regression against graph features, thus indicating that orientation, atom types and other structural features of the molecular graphs play a critical role in charge transfer.

To facilitate our understanding of the factors affecting electron transfer, we reduced the feature set to the top 10 uncorrelated features (≤0.7 correlation, see STAR Methods A.4), and we achieve a MAE of 0.361 ± 0.02 indicating that the majority of important information is captures by a small set of features (Figure 2B(ii)).

We used Shapley Additive Explanations (SHAP) [30], a game-theoretic framework that computes the contribution of each feature to the prediction of each sample (Figure 2B(ii)). Features with large absolute SHAP values have an overall large impact on prediction, whilst the sign of the SHAP values indicates a positive or negative effect of that feature on the prediction of the sample. The sum of absolute SHAP values across all samples for each feature allows us to assess their relative impact on the prediction task [30]. As seen in Figure 2B(ii), the top feature is the *variation in the edge betweenness centrality* (VEBC) of the molecular graph, and Figure 2B(iii) shows that an increase in VEBC correlates strongly with a decrease in *J*-coupling. Indeed, regressing against VEBC alone already gives a MAE of 1.032 ± 0.08, which is substantially lower than regressing against the distance between the molecules in the dimer. Edge betweenness is a graph property that measures the importance of an edge for mediating the shortest paths between nodes; hence a low variation in edge betweenness suggests a balanced spread of communication in the dimer, with no single atomic interaction acting as a critical funnel for the shortest paths. Another interesting graph feature in the top set is the *maximum rich club coefficient* (MRCC). As shown in Figure 2B(iii), a large MRCC is linked to large *J*-coupling. A large MRCC means that the hubs are well connected, and that the global connectivity is resilient to hub removal. In our molecular graphs, such robustness to the removal of particular atoms indicates the existence of alternative paths in the structure that may facilitate charge transfer between the molecules in the dimer. Further examination of the molecular features related to high charge transfer in helicenes lie beyond the aims of this work. However, further research could pose an associated classification task to identity graph features that lead to very high or very low *J*-couplings with the aim to guide the design of more efficient helicene compounds.

## 3 Discussion

We have introduced HCGA, a high-throughput computational graph feature analysis package that leverages the giant corpus of graph theory literature. The software package distills a large set of graph-theoretical algorithms making them easily accessible and simple to interpret for a given research problem. Beyond the massive feature extraction, HCGA includes a suite of statistical prediction tools for classification and regression to help researchers analyse their data sets. The highly comparative nature of HCGA provides a framework to identify individual features that play an important role in prediction and reveal scientific insights into their systems.

The use of network data in statistical learning is a current area of intense work[37]. For instance, researchers have turned to graph neural networks (GNNs) to learn structural features through message-passing and non-linear interactions[38, 39]. However, such methods lack the interpretability necessary to facilitate discovery science [40] and can be fraught with inaccuracies and inflated performance due to over-engineering [31]. The area closest in essence to HCGA is that of graph embeddings, in which the graph is reduced to a vector that aims to effectively incorporate the structural features [41]. However, the inherent choice of network properties that provide a ‘good’ vector representation of the graph is not known and may differ between scientific domains and the type of statistical learning task. HCGA thus circumvents this critical step in the embedding process through indiscriminate massive feature extraction.

We have illustrated the use of HCGA on a variety of examples drawn from different scientific domains performing different learning tasks (supervised classification, unsupervised clustering, regression). The framework is highly predictive and provides interpretable insights. HCGA has general utility and can be applied to a variety of fields where network data sets constitute the core of experimental outcomes, including applications to functional or structural connectivity networks in brains, revealing properties of computer networks that make them weak to outside attacks, the analysis of the structure of ecological networks and their fragility, the interdependencies in social and economic networks, and many other problems across scientific domains.

## 4 Acknowledgements

R.P., A.A, S.N.Y and M.B acknowledge funding through EPSRC award EP/N014529/1 supporting the EPSRC Centre for Mathematics of Precision Healthcare at Imperial. K.E.J. thanks the Royal Society for a University Research Fellowship and for an Enhancement Award 2017. We gratefully acknowledge advice and interpretation of results from Asher Mullokandov and Paul Expert.

## 5 Author contributions

R.P., A.A., M.B. conceived the framework. R.P., A.A., H.P., N.B. designed the methodology and wrote the software package. R.P., A.A., J.S. performed testing and benchmarking. J.S., K.E.J. developed the semiconductor data set and interpreted the results. R.P., A.A., J.S., S.N.Y, M.B. wrote the manuscript.

## A STAR METHODS

## A.1 Key resources table

| REAGENT or RESOURCE | SOURCE | IDENTIFIER |
| --- | --- | --- |
| Code repository | This manuscript | https://github.com/barahona-research-group/hcga |
| Code documentation | This manuscript | https://barahona-research-group.github.io/hcga/ |
| Code examples | This manuscript | https://github.com/barahona-research-group/hcga/tree/master/exam |

## A.2 Lead Contact

Further information and requests for resources should be directed to and will be fulfilled by the Lead Contact, Robert Peach.

## A.2.1 Data and Code Availability

The authors declare that the data supporting the findings of this study are available within the paper and its Supplemental Information files. The code is shared under the GNU General Public License v3.0.

## A.2.2 Benchmark data sets

Table 2 provides a high-level description of each data sets. The benchmark data sets were taken from https://ls11-www.cs.tu-dortmund.de/staff/morris/graphkerneldatasets.

## A.2.3 Neuronal morphology data set

The neuron data set is constructed from the morphological reconstructions used in the Blue Brain Project, freely available at http://microcircuits.epfl.ch/#/article/article_3_mph. A neuron contains a soma in its center, from which several types of branches emerge: axonal branches are carrying electrical signals away from the cells and basal or apical dendrites receive electrical signals from pre-synaptic cells, which will eventually trigger action potentials (or spikes) in the axon initial segment, near the soma. The shape of neurons is thus important in the electrical properties of the cells as well as how and where it connects to pre- and post-synaptic cells in a neuronal circuit. For this reason, an accurate and meaningful classification of shape (m-types) and electrical properties (e-types) of neurons is an active research topic in neuroscience, see for example [42]. Here, we will only consider morphological types, represented as graphs.

Most commonly, morphologies are classified by the reconstructors in morphological types, which depend on the cortical layer (from 1 to 6) from which they were extracted and a subjective interpretation of the shape of the cell. There are two broad classes of cells, pyramidal cells, or excitatory cells, which contain apical trees and interneurons, or inhibitory cells, which only have basal dendrites. We selected a subset of neuronal morphologies such that each m-type consists of at least 10 associated morphologies, consisting of 444 morphologies classified in 24 different m-types across 6 layers of the rat somatosensory cortex.

For each cell, we constructed a graph representation by extracting the neuron connectivity between sections using the python package MorphIO (https://github.com/BlueBrain/MorphIO), where a section is defined as a list of points along branches between two branching points. Each node of the connectivity graph represents a section, and an edge is present between two sections if they are connected by a branching point. This representation allows us to assign node attributes as the path length and the mean diameter of each section. The path length is defined from the three dimensional location of the points along the section as the sum of the length of each interval. In addition, we added a node label to represent the soma, as [1, 0] for the soma node, and [0, 1] for all other nodes.

To compute the pairwise classificaiton accuracy between each pair of m-types, we use a two-step procedure. We first used the entire feature set extracted with HCGA, secondly we chose a subset of the feature set containing the top 10 features that were less than 0.7 Pearson correlated between themselves. This drastically reduced set of features increased the classification accuracy on each pair of m-types (likely by reducing over-fitting to a large feature set).

Given the matrix of pairwise classification accuracies, we then applied a sigmoid function *f* (*x*) = 1*/*(1+*e*^−10(*x*−0.8)^) in order to make low and high accuracies more similar whilst retaining intermediate accuracies. This step was implemented simply to enable a more robust clustering of the similarity matrix when using the hierarchical clustering algorithm. The groupings of m-types we robust across different parameters for feature extraction (e.g. using all features or choosing only highly interpretable features to produce the similarity matrix) suggesting that the results reflect real clusters of m-types and are not an artefact of our feature choice.

## A.2.4 Organic semiconductors data set

The organic semiconductors (helicenes) data set was taken from a recent computational screening for high charge-carrier mobility study [6]. In the original study, a total of 1344 potential [6] helicene molecules were screened for their suitability as potential organic semiconductors. Here, we focus on the electronic transfer integral *J* (J-coupling). The electronic transfer integral *J* describes the hopping integral, i.e., the ease with which charge can hop from molecule A to B of the same type. This integral is strongly dependent upon the frontier orbitals of molecules and the spatial arrangement and orientation of molecules A and B, and its prediction in new molecules is currently an open challenge. For further details on the data set, see[6].

For each dimer (pair of molecules) at its minimum-energy separation *d*_*min*_, we computed the transfer integrals by projecting the computed orbitals of the dimer onto the unperturbed localised orbitals of the individual molecules [35, 43]. All transfer integrals were computed with B3LYP/6-31G(d) using Gaussian16[44].

In this data set only one spatial arrangement was used, the so-called translational-dimer motif. The electronic transfer integral is believed to exponentially decay with intermolecular distance, therefore the minimum energy distance *d*_*min*_ at which the *J* was computed is used as a control with which to compare HCGA.

For graph construction, each node represents an atom and edges corresponded to atom interactions. To define atomistic interactions we used the relative spatial distances between them; if the separation of two atoms was lower than the sum of their Van-der-Waals radii plus 2Å then we built a edge between the two atoms. The weight of the edge was simply the inverse of the Euclidean distance between the two atoms (calculated given their xyz coordinates). Each node was also assigned features based on the atom type using a one-hot encoded vector.

## A.3 Method details

## A.3.1 Software

Our entire pipeline is self-contained within a Python environment and only requires the user to input their data set in the appropriate format (format described in Section A.4.2).

The pipeline can be implemented via two routes. The first is to use a shell script that allows the continuous execution of the entire pipeline. To test the benchmarks, we have included a script that automatically downloads and pre-processes the necessary data set. Python 3 scripts then execute the feature extraction which relies on the NumPy, Pandas, NetworkX and SciPy environments. The post-processing and statistical analysis steps are implemented using the sklearn and SHAP environments.

To enable continued preservation of the Python software package we have designed a modular and robust framework. Features can be added and removed easily and external users are able to make pull-requests to the main repository on github.

## A.3.2 Run times

Computational efficiency is a key problem in deriving graph statistics, as, for example, features based on k-components or node-connectivity may take on order of minutes to compute on a high-end computer. To mitigate these issues we offer the user two options; (1) a time-out option where each feature is given a time-limit (default 10 seconds) before NaN is returned, and (2) a feature speed ranking (’slow’,’medium’,’fast’) allowing the computation of only a subset of faster features. Feature extraction for the benchmark data-sets is normally completed on the order minutes to hours on a local computer. Most features are based on the python package networkx, but faster implementations of some features may be used. To allow for the use of other network packages to extract features, the internal graph representation is generic, and only at the level of feature extraction a specific representation is used, such as with networkx graph for example.

## A.3.3 Feature Interpretability

A key aspect of HCGA is the feature interpretability that provides researchers key insights. However, even classical graph theoretic measures can be difficult to interpret in respect to most systems. Therefore, we have manually assigned an interpretability ranking (from most interpretable 5 to least interpretable 1) to each feature allowing users to implement statistical analysis with a chosen level of interpretability.

## A.3.4 Feature filtration and normalization

Depending on the data set, some features may not be computable for particular graphs. These values need to be removed prior to statistical analyses. Any features with infinity or not a number (NaN) values and features with zero variance across the data set are removed from the feature matrix, resulting in a reduced feature matrix. The quantity of removed features is dependent on the data set, and often small if the data set is composed of relatively similar graphs. In some cases, particular graphs can result in the removal of a large set of features, therefore it is advised to explore anomalous samples. Finally, each remaining feature is individually standardised to have zero mean and a unit variance in order to guarantee stable convergence during statistical analysis.

## A.3.5 Graph classification

Graph classification is the process of predicting the class for each graph by mapping a function from the feature matrix to a vector of discrete output variables. To use this functionality the user must have input a set of class labels, one for each sample. The HCGA package allows the user to use any classification algorithm, however, the default procedure (and the procedure used within this paper) is to use the XGBoost classification algorithm with default parameters[45]. XGBoost is a decision-tree-based ensemble Machine Learning algorithm that uses a gradient boosting framework, considered the optimal approach for small-to-medium structured/tabular data.

The classification procedure uses a 10-fold stratified cross-validation with no hyper-parameter optimisation. Our choice to not use hyper-parameter optimisation was to provide a use-case in which a non-machine learning expert was using our software package. With careful tuning, a researcher will be capable of improving their results. Quoted results in this paper are the average across the 10-folds, this was used to prevent over-fitting leading to optimistic performance estimates.

## A.3.6 Graph regression

Graph regression is the process of predicting a continuous variable for each graph by mapping a function from the feature matrix to a vector of continuous output variables. To use this functionality the user must have input a continuous output variable for each sample. The HCGA package allows the user to use any regression algorithm, however, the default procedure (and the procedure used within this paper) is to use the XGBoost regressor algorithm with default parameters[45].

Similarly to classification, no hyper-parameter optimisation is implemented and results are averaged across a 10-fold stratified cross validation.

## A.4 Reduction of feature set

Due to the large feature set, the training set may be overfitted, resulting in a decrease in accuracy on the test set. To remedy to this problem, we implemented a simple procedure for reducing the size of the feature set. From the first classification/regression step with all the features, we computed the SHAP importance value of each features as well as the Pearson correlations between all of them. We then chose the *n* most important features which are correlated less than a value *c*. These two numbers, *n* and *c* are the only parameter the user can modify to reduce the feature set. We have set as default *n* = 100 and *c* = 0.8 for the results shown in Tables 3 and 3, however, these values can be optimised with a validation set (see starred HCGA results in Tables 3 and 3).

## A.4.1 Feature Importance

HCGA provides implements current methods for investigating the contribution or importance of individual features towards the classification or regression problem. Specifically, we compute top features using the Shapley Additive Explanations (SHAP) framework which computes optimal explanations based on game theory through local feature interaction effects[30]. We compute the SHAP values of every feature for each fold in our cross-validation procedure and return an average across the folds. We output a SHAP value for each feature given its ability to separate each class individually. We also average SHAP values for each class to return an overall feature importance measure across the entire set of classes.

## A.4.2 Inputs

The benchmarks can be automatically downloaded without any need to provide custom input into HCGA. For custom data we have provided a number of example notebooks to aid the user in their application. We have built a generic Graph object which allows the user to pass their graph to HCGA and be automatically converted to the appropriate graph representation used by the feature extraction module. The user should provide a graph, any node features and a graph label (or continuous variable).

## A.4.3 Outputs

The primary outputs from HCGA are the computed features for each input graph. The list of master operations are detailed in Table 1. The secondary outputs of HCGA are a result of the statistical analysis module. This includes a comma-separated value file, that details the importance of each feature (SHAP value) towards the statistical prediction task (classification or regression), and a results report which includes a series of plots to facilitate user insights into their data. The plots include:

1. *Bar plot detailing the mean absolute SHAP value for the top features.* A larger value indicates that the feature had a larger impact on the prediction task.
2. *A sample expanded feature summary*. Detailing the impact of each feature on individual samples.
3. *A heatmap of absolute correlation coefficient between features.* The heatmap is ordered by the feature type. Green dots along the diagonal indicate the top features.
4. *A heatmap of absolute correlation coefficient between features.* The heatmap is clustered according to Euclidean distance within the space of correlations (clustering applied using a Ward linkage algorithm). Green dots along the diagonal indicate the top features.
5. *Plots of individual features.* In classification tasks, these appear as violin plots for each class, and for regression tasks these appear as a scatter plot of dependent versus independent variable.
6. *Feature summaries for each top feature.* A violin plot is displayed for each feature and representative sample networks are displayed alongside representing different points in the feature distribution.

## B Graph-theoretical features

The core functionality of HCGA is to exhaustively compute all graph theoretic features. Methods for graph analysis can take a variety of forms, from simple summations to complex convolutions. In HCGA, we have implemented each such method as an algorithm: an operation that summarises an input graph with a single real number. The real-valued summary values are collected into a feature vector representation of a graph *G*_*i*_. Repeating this process for each graph *G*_*i*_ in a collection of graphs 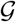 outputs a feature matrix. Below we describe at a high-level the different types of features that are computed and in Table 1 we detail the main feature classes that are implemented to date.

## B.1 Graph level features

There exist a number of graph theoretic features that output a single value that describes the network. These values can therefore be implemented directly as features that describe a given graph. For example, the number of nodes in a network is a simplistic description but offers a single valued feature of the network. The small world coefficient, computed as the ratio of average clustering coefficient to average shortest paths, is also a single valued descriptor.

## B.2 Summary statistics.

Many graph-theoretic algorithms, such as measures of node centrality, output many values which then need to be reduced to a few representative quantities that provide a single valued descriptor of the graph. We implemented two approaches to summarise vector (or higher dimensional) outputs from graph-theoretic algorithms. If the output is a distribution, we used a summary statistics function that implements basic statistical quantities such as mean, variance, or kurtosis among others. If the output is a series of sets (e.g. the output from community detection would give sets of nodes assigned to communities) then we implement a series of clustering quality algorithms such as modularity or coverage.

## B.3 Features from node/edge attributes.

In addition to features solely based on the topology of the graphs, some data sets contain relevant node or edge attributes that may provide a more detailed description of the modelled system. We can compute statistics that summarise the node or edge attributes as additional descriptors of the network. For example, given a single node attribute such as the atom type in a molecular graph, we can compute the sum of each atom type. Taking inspiration from graph diffusion methods [46, 10], we also compute features that summarise the transformation of the node attributes by the graph topology via message passing across the adjacency matrix. In addition, there may be a number of possible extensions involving both edge and node features or even other representations of graphs, such as for example multilayer networks[47], which we leave for future works.

## C Extended Results

## C.1 Benchmark data sets

The complete set of classification results are detailed in Tables 3 and 4 against popular deep-learning methodologies and Kernel algorithms using the results reported under Fair Comparison[31].

All benchmarks were run with the same HCGA conditions: timeout for feature extraction of 30 seconds with all possible features; XGBoost classifier with default (unoptimised) parameters with 10-fold stratified splits; feature set used for classification is the top *r* = 100 features with a maximum correlation of *c* = 0.8. The accuracies so obtained are reported in Tables 3 and 4.

We also ran the benchmarks under a slightly modified HCGA∗ classification: timeout for feature extraction of 30 seconds with all possible features; XGBoost classifier with default (unoptimised) parameters with 10-fold stratified splits; feature set used for classification obtained by optimising against a validation set (shown for selected data sets at the bottom of Tables 3 and 4) using a manual and coarse grid-search to obtain a set of *r* features with maximum correlation *c*, e.g.,

- Enzymes: *r* = 125, *c* = 0.99
- Proteins: *r* = 50, *c* = 0.90
- D&D: *r* = 75, *c* = 0.90
- NCI1: *r* = 75, *c* = 0.95
- Mutag: *r* = 50, *c* = 0.95
- Collab: *r* = 100, *c* = 0.99
- IMDB-B: *r* = 75, *c* = 0.99
- IMDB-M: *r* = 25, *c* = 0.99
- REDDIT-B: *r* = 100, *c* = 0.95
- REDDIT-5K: *r* = 50, *c* = 0.95

## C.2 PROTEINS data set insights

As an example of the feature interpretability that HCGA offers, we have performed a more detailed review of the proteins data set. The proteins data set is comprised of 1178 high-resolution protein graphs belonging to two classes: enzymes (59% samples) and non-enzymes (41% samples)[53]. Each protein graph has amino acids as nodes and edges defined by the bonds between amino acids. In addition, each node includes information about whether the amino acid is a member of a beta sheet, an alpha helix or a turn[54]. The prediction of protein function is a difficult task given that no chain in any protein aligns to any other chain in the data set. In the original paper[53], node attributes were crucial for improving the classification accuracy of a SVM. Here, node attributes remain important, but are not crucial for HCGA.

Figure 3 displays the top features that best differentiate the two classes in the proteins data set. We find a mixture of structural features and node features. For instance, the top feature is the total number of cliques in the graph and we find that the enzymatic class exhibits a larger total number of cliques relative to the non-enzymatic class. A clique is a set of nodes such that every two nodes in the set are adjacent; hence cliques represent compact structural regions with large number of interactions among the residues, indicating structurally stable regions of the protein [55, 4] that are required for enzymatic activity. Other structural features include the mean rich club and the standard deviation of the page rank, both commonly used graph theoretical methods for analysis across a broad range of scientific domains. Looking at node features, we find that the standard deviation of the first and second node features both appear as important (‘node_feature0_std’ ranked 13th, ‘node_feature1_std’ ranked 6th, out of all extracted features). Node features 0 and 1 indicate that a node was found inside an *α*-helix or *β*-sheet respectively. The standard deviation is computed across the vector of node membership for each feature and a larger value here is indicative of a larger fraction of secondary structure occupancy. We find that the standard deviation of both node features (0 and 1) was higher for the enzymatic class, suggesting a larger *α*-helix and *β*-sheet occupancy for enzymatic proteins, further agreeing with the structural feature (total number of cliques) that indicates the increased number of structural regions within enzymatic proteins. The results here highlight the importance of both structural features and node features in providing the best prediction.

Figure 4 shows the clustered correlation matrix of the top features for the proteins data set. Clustered features share a similar profile in the data set and also reflect an overlap in the definition of the graph properties, i.e. features based on rich club coefficients are clustered together. This plot allows the researcher to identify redundant features so as to select sets of interpretable, yet independent, features to best describe the system of interest.

## Footnotes

2 The HCGA python package is available at https://github.com/barahona-research-group/hcga.

## Notes

https://github.com/barahona-research-group/hcga

### Competing Interest Statement

The authors have declared no competing interest.

### Summary of Updates

The revision has been made to make a spelling mistake in an authors names and also to extend the literature review within the introduction.

